# PAH-former: Transfer Learning for Efficient Discovery of Pulmonary Arterial Hypertension-Associated Genes

**DOI:** 10.1101/2025.06.16.660027

**Authors:** Toshinaru Kawakami, Sosuke Hosokawa, Masamichi Ito, Atsumasa Kurozumi, Ryohei Tanaka, Shun Minatsuki, Junichi Ishida, Takayuki Isagawa, Satoshi Kodera, Norihiko Takeda

## Abstract

Single-cell RNA sequencing (scRNA-seq) of patient samples holds promise for understanding disease mechanisms, but faces the challenge of excessive cost and effort in acquisition, processing, and data analysis, making it essential to leverage existing data. Pulmonary artery hypertension (PAH) is a refractory disease characterized by pulmonary vascular remodeling, and access to patient specimens is limited due to difficulties in tissue collection. In this study, we employed transfer learning with Geneformer, a deep learning algorithm pre-trained with scRNA-seq datasets and fine-tuned it with public PAH lung tissue data to identify the disease-relevant genes. The resulting algorithm, which we named PAH- former, demonstrated that its prediction accuracy varied significantly depending on the dataset used for fine-tuning. PAH-former enabled us to perform *in silico* perturbation analysis and identified PAH related genes. Loss-of-function PAH related genes in human pulmonary artery endothelial cells increased the expression of *SOX18*, a signature gene of PAH. This integration of artificial intelligence and biological experiments can significantly advance our understanding of molecular mechanisms of PAH.

## Introduction

PAH is a severe and complex disease marked by high pulmonary arterial pressure, causing right heart failure and death ^1^. Pulmonary vascular remodeling is a common and critical pathogenic feature of PAH. This process, considered largely irreversible, involves the dysfunction of endothelial cells (ECs), vascular smooth muscle cells, and fibroblasts, as well as the participation of immune cells ^2^. However, effective treatments are lacking due to its rarity and the absence of accurate *in vivo* models mimicking human disease. Identifying drug targets is thus challenging, often requiring lengthy knockout animal validation. Moreover, understanding PAH fully requires cell-specific analysis to clarify each cell type’s molecular roles. Therefore, innovative methods are essential to overcome these hurdles and discover effective PAH therapy targets. Notably, *SRY-Box Transcription Factor 18 (SOX18)*, a key transcription factor, has been identified as a signature gene whose expression is significantly upregulated in PAH, particularly in endothelial cells. Its involvement in angiogenesis and endothelial function highlights its importance in PAH pathogenesis, making it a valuable indicator of disease-like cellular states ^3,4^.

scRNA-seq technologies have emerged as powerful tools for dissecting complex diseases, offering unprecedented resolution to identify key genes and pathways within individual cell populations. However, analyzing these vast datasets and translating them into useful insights is not straightforward. One major hurdle lies in the absence of robust algorithms for prioritizing candidate disease genes from the extensive lists generated by differential gene expression (DEG) analysis. The selection process often relies on subjective criteria and the specific databases employed, potentially introducing bias and limiting reproducibility.

Furthermore, DEG analysis alone does not guarantee the identified genes are causally implicated in the disease pathogenesis. In many rare diseases like PAH, obtaining large cohorts of human samples for comprehensive validation remains a significant bottleneck due to the severity of the disease and the technical difficulties of tissue collection.

Geneformer ^5^ is a foundational transformer model pre-trained on a large-scale corpus of single-cell transcriptomes to enable context-aware predictions in network biology. It was originally trained on approximately 30 million single-cell transcriptomes in June 2021, and later expanded to about 95 million transcriptomes in April 2024. Some researchers have already applied Geneformer to a variety of downstream tasks. Mellors *et al.* fine-tuned it with bulk tumor gene expression data and proposed a novel transformer model predicting the tissue of origin for cancers ^6^. Chen et al developed a model to predict tumor-restricting factors in the colorectal tumor microenvironment using a cancer-tuned Geneformer and *in silico* treatment analysis ^6,7^. Wang *et al.* constructed context-specific brain gene regulatory networks. They fine-tuned Geneformer with brain single nucleus RNA-seq data and conducted *in silico* gene perturbation studies. They further applied these networks to the study of autism spectrum disorder ^8^. Geneformer, with its ability to learn complex gene expression patterns and relationships from massive transcriptomic data in various conditions, offers an unbiased and robust approach to identify disease associated genes beyond DEG analysis. Although applications in understanding clonal pathologies such as cancer have been reported, there is no precedent for its use in elucidating complex systemic conditions like cardiovascular diseases. This is due to the lack of sufficient published data sets for fine-tuning in this area and the absence of studies that have experimentally validated the application of Geneformer.

In this study, to overcome these challenges and improve scRNA-seq data analysis quality, we built a novel platform based on Geneformer (PAH-former) and trained it based on public data of PAH. We also tested the effectiveness of addition of datasets in improving prediction accuracy and validated the established models by *in vitro* experiments. Our approach not only avoids the limitations of traditional DEG analysis based methods but also demonstrates the broader applicability of Geneformer based fine-tuning as a powerful strategy for identifying disease associated genes. This study leverages our novel platform to identify and validate new disease associated genes of PAH, promising to advance our understanding of cell specific disease mechanisms and pave the way for novel therapeutic strategies.

## Results

We conducted fine-tuning of Geneformer by public scRNA-seq analysis data to create “PAH-former”, which can efficiently detect PAH associated genes (Fig. 1A). PAH-former was trained using publicly available idiopathic pulmonary arterial hypertension (IPAH) datasets and can be utilized for various downstream analyses, such as cell type prediction and *in silico* perturbation. Single-cell data of IPAH is very limited and we primarily utilized data from GSE169471. First, we re-performed clustering and cell type annotation using the raw data from GSE169471 with CellTypist v2.0 ^9^. As a result, we achieved clustering and cell type annotation comparable to the t-SNE presented in the original paper ^3^ (Fig. 1B). We proceeded to map the cells, distinguishing between PAH and healthy groups. Our analysis revealed that each cluster contained cells from both control and PAH, aligning with the results presented in the original paper (Fig 1C). As will be discussed later, the original paper demonstrated upregulated expression of the transcription factor *SOX18* in endothelial cells of the PAH group. When we mapped the expression levels of *SOX18,* we similarly observed its selective expression in endothelial cells (Fig 1D). These results collectively indicate the accuracy of our clustering and annotation.

**Figure 1.**
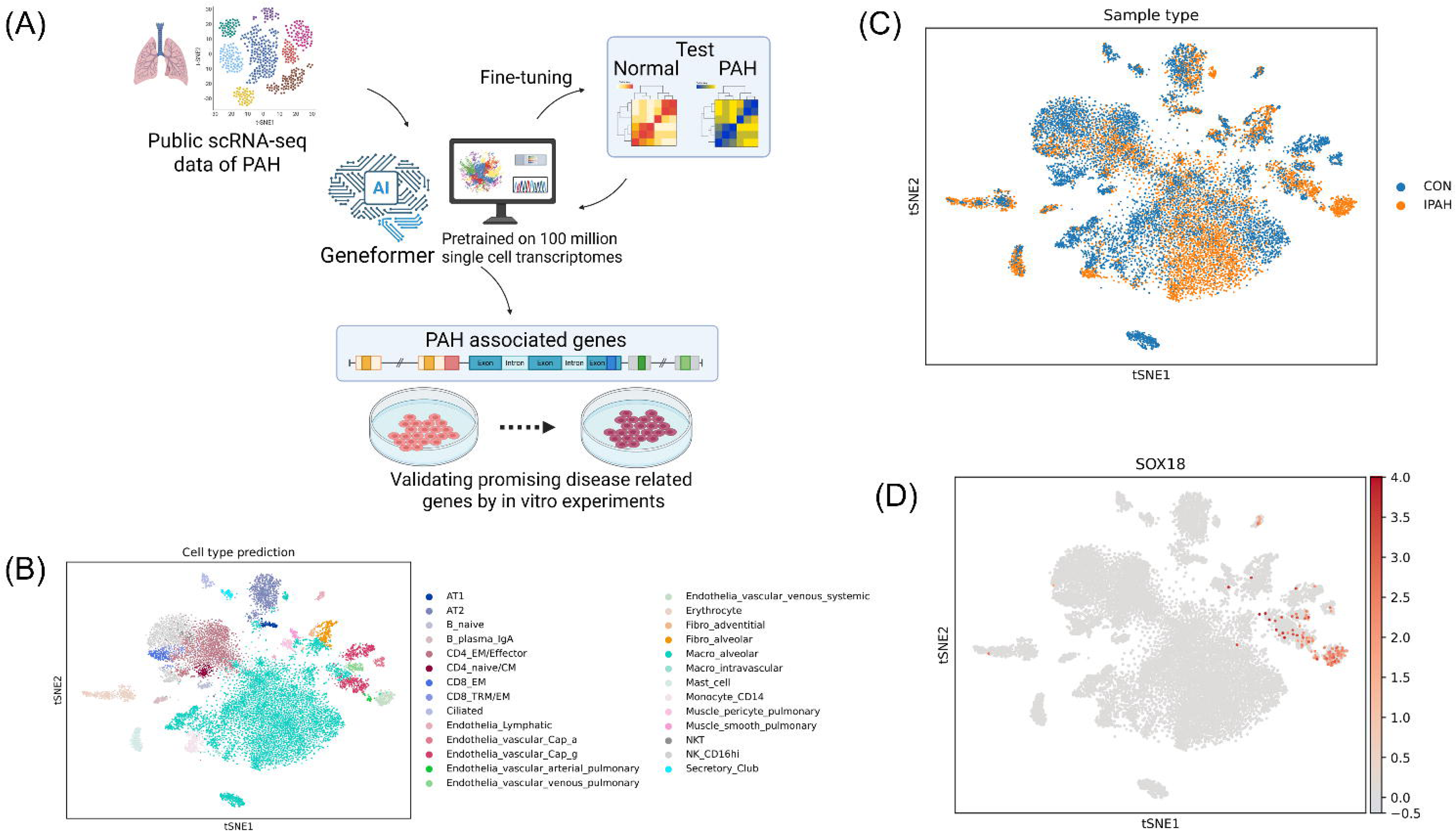
Public single cell RNA-seq data of pulmonary artery hypertension lung and its reanalysis. (A) Schematic of the PAH-former development pipeline. We created a new AI tool that understands the gene expression network of pulmonary arterial hypertension (PAH) by fint- tuning Geneformer, a transfer learning tool that has been trained on 1 billion single-cell analysis data, with publicly available PAH single-cell analysis data. The aim is to extract the genes that are involved in PAH. scRNA-seq,single-cell RNA sequencing; t-SNE, t-distributed stochastic neighbor embedding; PAH, pulmonary arterial hypertension. (B) t-SNE plot showing cell type prediction. Cell type annotation of the GSE169471 data was performed using CellTypist v2.0. t-SNE, t-distributed stochastic neighbor embedding. (C) t-SNE plot showing sample type distribution. CON,control; IPAH, idiopathic pulmonary artery hypertension; t-SNE, t-distributed stochastic neighbor embedding. (D) t-SNE plot visualizing *SOX18* expression levels. t-SNE, t-distributed stochastic neighbor embedding.

### Training Dataset Setup

According to Geneformer’s *in silico* perturbation of cardiomyocytes in the original paper, the training data included 93,589 cardiomyocytes (non-failing, n = 9; hypertrophic, n = 11; dilated, n = 9); the test data consisted of 39,006 cardiomyocytes (non-failing, n = 4; hypertrophic, n = 4; dilated, n = 2) ^5^. However, publicly available IPAH data is very limited, and its quality also varies, making it potentially difficult to secure a sufficient amount for fine-tuning of Genformer for IPAH. While using large datasets for fine-tuning carries a risk of overfitting, it has been reported that fine-tuning can be performed efficiently even with small datasets ^10,11^.

We compared three distinct training approaches and found that the inclusion of large control data from Human Lung Cell Atlas (HLCA) significantly enhanced the model’s accuracy and F1 score in cell classification. Figure 2A presents a table outlining the datasets used to train the three fine-tuning models (model A, model B, and model C) evaluated in this study. Model A was fine-tuned using only data from GSE169471, which training data included 6 samples, (control = 4 samples, 5106 cells, IPAH = 2 samples, 6514 cells). Model A exhibited a very poor performance, achieved an accuracy of 0.523 and an F1 score of 0.522 (Fig. 2B, C).

**Figure 2 .**
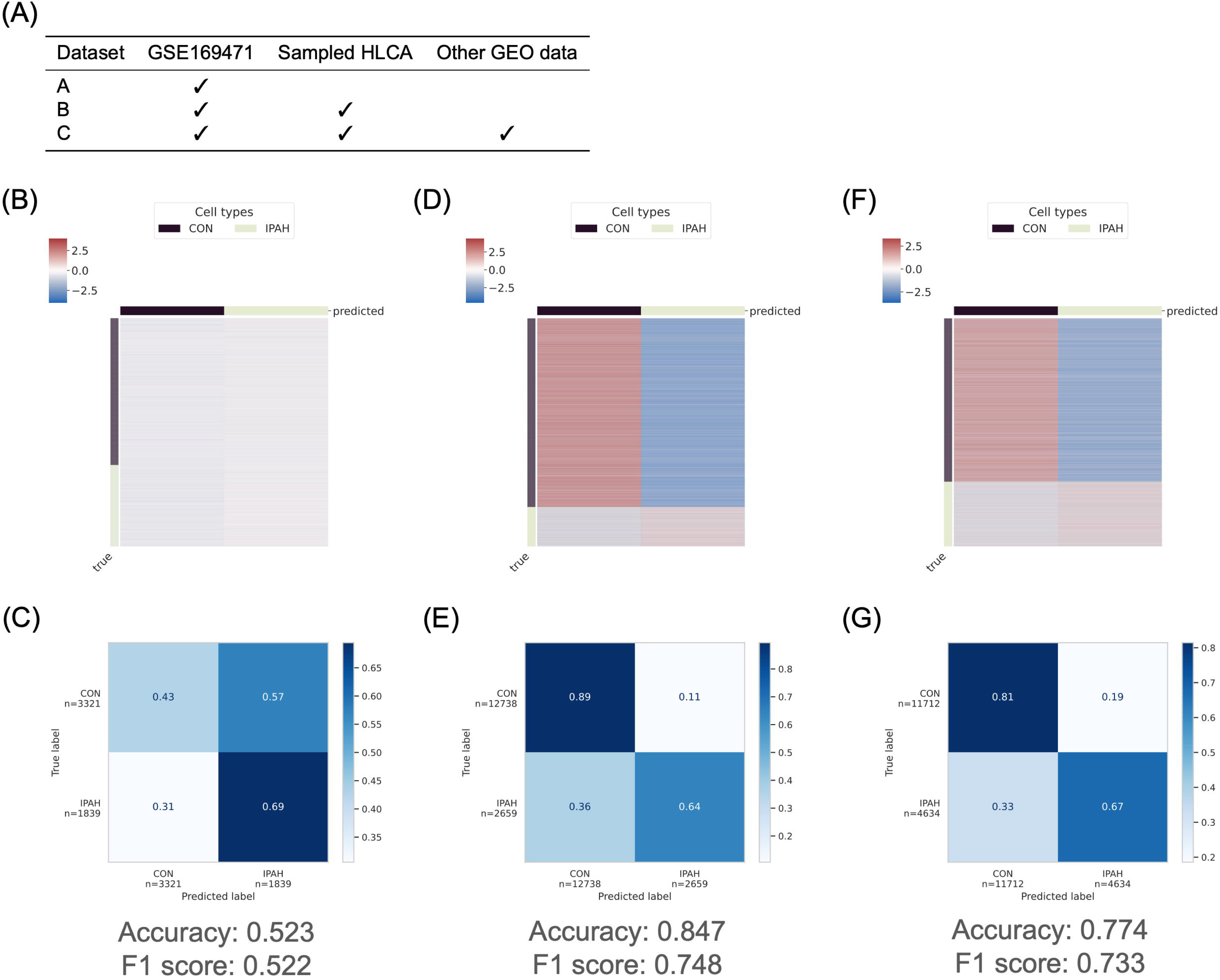
Dataset selection for the fine-tuning of Geneformer. (A) Datasets used for Geneformer model training. This table outlines the composition of training data for each model. HLCA, human Lung Cell Atlas. (B, C) Prediction likelihood heatmap for cell classification and confusion matrix of Model A. (D, E) Prediction likelihood heatmap for cell classification and confusion matrix of Model B. (F, G) Prediction likelihood heatmap for cell classification and confusion matrix of Model C. CON,control; IPAH, idiopathic pulmonary artery hypertension.

Model B augmented the GSE169471 dataset with a large quantity of healthy control cells sampled from HLCA (102 samples, randomly sampled 10,000 cells). Model B demonstrated a dramatic improvement in performance, achieving an impressive accuracy of 0.847 and an F1 score of 0.748. This substantial increase highlights the effectiveness of augmenting training data with large quantities of relevant control cells to better define the baseline healthy state (Fig. 2D, E). Model C represents an extended training approach, incorporating additional IPAH data from other GEO datasets (GSE210248, GSE185479; total 8 samples, randomly sampled 6,000 cells) along with GSE169471 and sampled HLCA data. Model C showed an accuracy of 0.774 and an F1 score of 0.733 (Fig. F, G). While superior to Model A, its performance was slightly lower than that of Model B. This observation suggests that for this specific task, the quality of control data from HLCA had a more significant positive impact than simply increasing the number of IPAH data.

### *In silico* perturbation analysis

To investigate the impact of specific genes on cell state in PAH, we performed in silico deletion and overexpression analyses using Geneformer fine-tuned by PAH scRNA-seq data (GSE169471) (Fig. 3A). To train Geneformer on scRNA-seq data, we performed rank value encoding, a method of ranking all genes in descending order based on their expression levels. *In silico* deletion refers to removing a specific gene from a list, while in silico overexpression refers to setting the rank of that gene to the first position. We conducted *in silico* perturbation analysis to investigate the directional shifts in cell state (cell embedding) following the deletion or overexpression of individual genes. Specifically, we assessed whether the cell state shifted from a healthy phenotype towards a PAH phenotype, or conversely, from the PAH phenotype towards the healthy phenotype. This analysis encompassed four distinct perturbation scenarios. As a result, we generated lists of putative disease associated genes for each of the four scenarios, which included genes previously implicated in the disease. Herein, we report the top 40 candidate genes identified for each scenario, along with the corresponding results of GO analysis of genes in each list (Fig. 3B- E).

**Figure 3.**
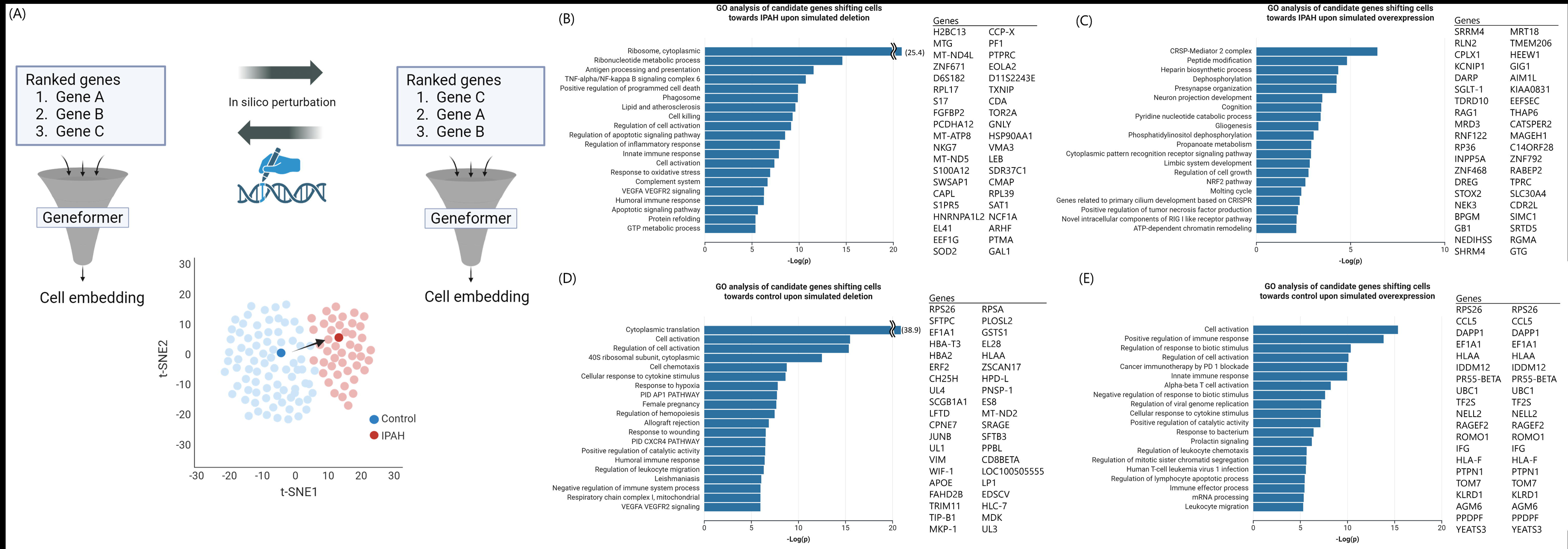
*In silico* perturbation using PAH-former and extraction of disease-related genes. (A) The workflow for *in silico* perturbation analysis using the fine-tuned Geneformer model (PAH-former). *In silico* manipulation (deletion or overexpression) results in shifts in cell embedding (representing cell state). IPAH, idiopathic pulmonary artery hypertension. (B-E) Gene Ontology (GO) analysis of candidate genes identified by *in silico* perturbation analysis by PAH-former and top 40 genes that shift cell embedding most for each of the four directions. (B) Gene Ontology (GO) analysis for candidate genes whose *in silico* deletion shifts the cell state towards IPAH. (C) Gene Ontology (GO) analysis for candidate genes whose *in silico* overexpression shifts the cell state towards control. (D) Gene Ontology (GO) analysis for candidate genes whose in silico overexpression shifts the cell state towards IPAH. (E) Gene Ontology (GO) analysis for candidate genes whose in silico deletion shifts the cell state towards control.

The number of candidate genes extracted by the fine-tuned Geneformer that shifts cell embedding from control to PAH state after in silico deletion were 134 (Supplementary Table X). Among the identified genes, while some were previously reported, the majority of them were novel. Previously reported genes included *HMGB2* (high-mobility group box 2).

HMGB2 is upregulated in PAH, and it is mentioned as a significant contributor to the pathogenesis of pulmonary hypertension by promoting inflammation and vascular remodeling ^12^. In addition, *SOD2* (superoxide dismutase 2) was also on the gene list, and its tissue specific, epigenetic downregulation initiates and sustains PAH by impairing redox signaling and promoting a proliferative, apoptosis-resistant pulmonary artery smooth muscle cell phenotype ^13^. This result is consistent with the cell embedding shifting from control to PAH state through *in silico* deletion. As described above, *in silico* deletion following fine-tuned Geneformer successfully identified a range of PAH related genes, including previously known genes. Notably, many genes in this list were not previously linked to pulmonary hypertension, suggesting them as novel candidates for exploring disease related molecular functions and pathways. There was minimal overlap between the gene set identified by our fine-tuned Geneformer and that derived from the DEG analysis in the original article, with only two genes being common to both lists (Fig 4A, 4B).

**Figure 4.**
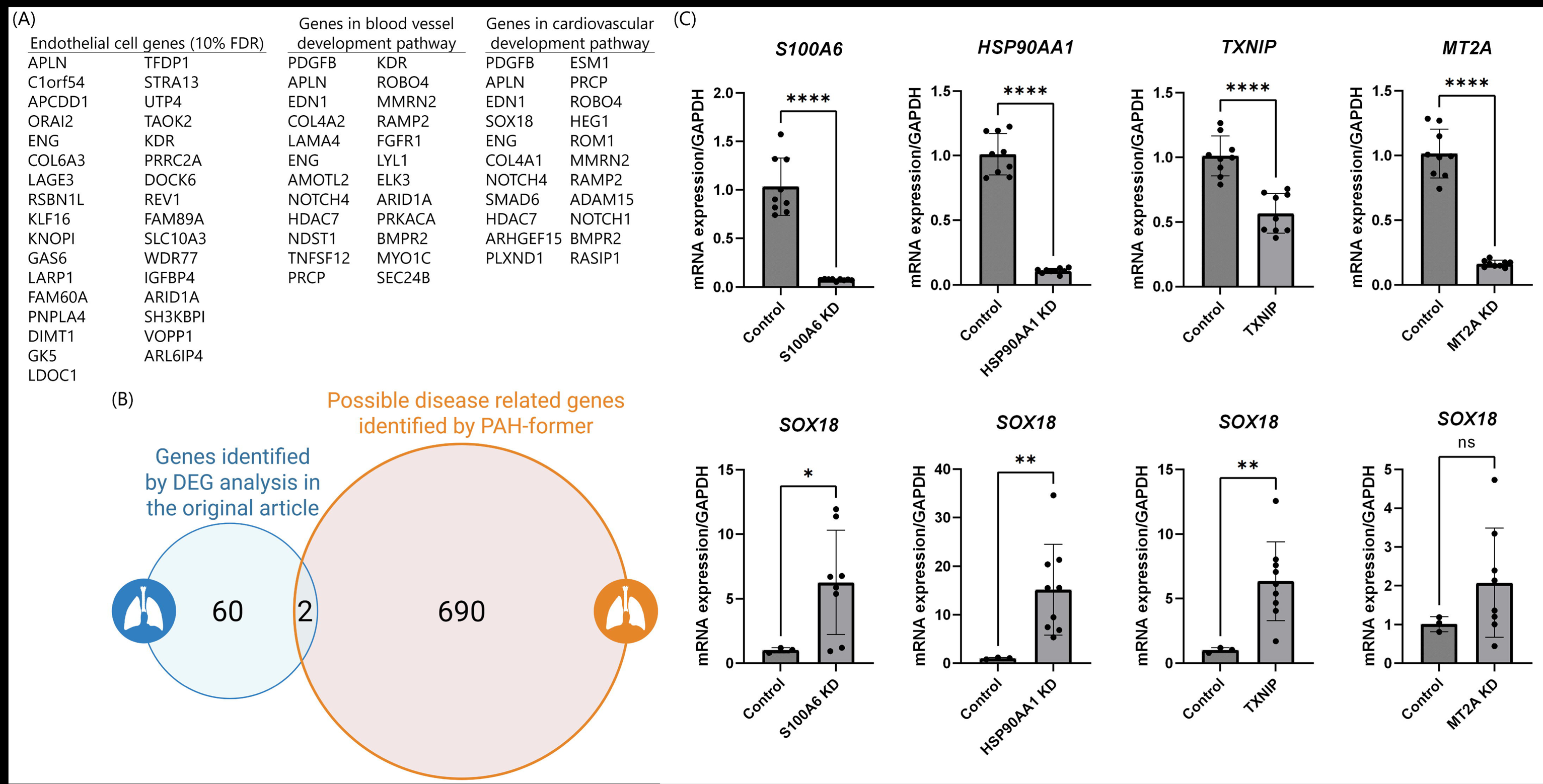
*In Silico* perturbation in pulmonary endothelial cells using PAH-former and *in vitro* validation. (A) Table showing partial lists of genes identified in endothelial cells by Differential Gene Expression (DEG) analysis (FDR < 10%) in the original article and genes enriched in the GO (gene ontology) pathways of blood vessel development pathway and cardiovascular development pathway. (B) Venn diagram comparing the number of candidate genes identified by Differential Gene Expression (DEG) analysis in the original article and the number of disease-associated candidate genes identified by PAH-former. PAH, pulmonary arterial hypertension. (C) Knockdown experiments of selected candidate genes (*S100A6, HSP90AA1, TXNIP,* and *MT2A*) using siRNA in Human Pulmonary Artery Endothelial Cells (HPAECs). Top panels show the knockdown efficiency of each target gene relative to control siRNA, displaying relative mRNA expression levels at 48 hours and indicating successful knockdown (data normalized to *GAPDH* expression, n = 9). Bottom panels show the relative mRNA expression levels of *SOX18* at 48 hours after knockdown of each candidate gene compared to control siRNA (data normalized to *GAPDH* expression, n = 9). Graphs are presented as mean ± standard error of the mean (SEM). Statistical significance is indicated as: * *p* < 0.05, ** *p* < 0.01, **** *p* < 0.0001, ns: not significant.

### Enrichment analysis

Enrichment analysis of 134 candidate genes that shifted cell embedding from control to PAH state after in silico deletion highlighted the enrichment in TNF-α/NF-κB signaling, regulation of inflammatory response, oxidative stress response, and VEGFA/VEGFR2 signaling pathways. These findings indicate that our approach successfully identified pathways associated with PAH using PAH-former.

### Target Gene Knockdown

Among the genes listed as PAH-related, we identified four genes (*S100 Calcium Binding Protein A6 [S100A6], Heat Shock Protein 90 Alpha Family Class A Member 1 [HSP90AA1], Thioredoxin Interacting Protein [TXNIP],* and *Metallothionein 2A [MT2A]*) that have not been previously reported in association with PAH.

In the original paper on which our dataset is based, several transcription factors, particularly *SOX18*, were found to be upregulated and implicated in regulating the PAH endothelial cell transcriptome. *SOX18* is also reported to increase the expression of nicotinamide phosphoribosyltransferase (NAMPT) and is shown to be involved in the pathophysiology of PAH via NAMPT ^4^. Therefore, in order to determine whether the cellular state was approaching a PAH-like state after knockdown of candidate genes found by PAH-former, we chose to compare mRNA expression of *SOX18*.

We performed a knockdown experiment of the four candidate genes using RNA interference with human pulmonary arterial endothelial cells (HPAECs). For each gene targeted, the knockdown was successful. Interestingly, the knockdown of three of these four genes resulted in a significant increase in *SOX18* expression 48 hours after siRNA transfection (Fig. 4C), ensuring the validity of the candidates extracted by PAH-former.

## Discussion

It has been demonstrated that disruption of certain genes can lead to the development of phenotypes indicative of PAH. These genes include *Prolyl Hydroxylase Domain-Containing Protein 2 (PHD2), GATA Binding Protein 6 (GATA-6), Bone Morphogenetic Protein Receptor Type 2 (BMPR2), Tet Methylcytosine Dioxygenase 2 (TET2), NLR Family CARD Domain Containing 3 (NLRC3),* and *AMP-Activated Protein Kinase (AMPK)* ^14–19^. *BMPR2* mutations, for example, cause PAH through the mechanisms such as endothelial dysfunction, smooth muscle cell abnormalities, mitochondrial dysfunction, and inflammatory responses ^20–22^.

We reanalysed the open-source single cell RNA-seq data of IPAH ^3^. Based on the findings by Saygin *et al.*, the expression level of the transcription factor *SOX18* was significantly upregulated in endothelial cells from IPAH patients. *SOX18* is known to be involved in angiogenesis and the regulation of endothelial barrier function, suggesting its critical role in IPAH pathogenesis ^23–25^. Sun *et al.* elucidated the complex regulatory network governing *NAMPT* expression in PAH and found that SOX18 plays an important role alongside STAT5, SOX17, and HIF-2α. SOX18, while exhibiting context-dependent effects, is a key regulator of VEGF-induced *NAMPT* promoter activity, and was shown to act in a manner that is opposite to SOX17 ^4^ . Therefore, we decided to use *SOX18* mRNA expression to assess whether the cellular state was approaching an IPAH-like state following knockdown of candidate genes identified by PAH-former.

The candidate genes were *S100A6*, *HSP90AA1*, *TXNIP* and *MT2A*. Although these genes have not been previously reported in association with PAH, their known biological functions offer plausible connections to disease mechanisms.For instance, *S100A6* (also called *Calcyclin, Cacy*) is a Ca2+-binding protein involved in cell proliferation and stress, with previous research suggesting its role in regulating antiproliferative pathways and potential interactions with proteins like Calcyclin-Binding Protein and Siah-1 Interacting Protein (CacyBP/SIP) in PAH models, hypothetically impacting vascular remodeling ^26–28^. *HSP90AA1* plays a crucial role in maintenance of endothelial nitric-oxide synthase (eNOS) dimer stability in pulmonary arterial endothelial cells and is an upregulated immune-related gene in PAH, implying its involvement in endothelial dysfunction and inflammatory processes ^29,30^. *TXNIP* mediates oxidative stress by inhibiting thioredoxin activity, a system linked to PAH progression, suggesting that its knockdown could enhance pro-PAH conditions ^31–34^.

Lastly, MT2A, a member of the metallothionein family, possesses antioxidant properties and is elevated in PAH patients, indicating its potential role as a biomarker and a defense against oxidative stress; its deletion might compromise this protection, leading to PAH-like pathology35.

Although these genes have entirely distinct functions, the common upregulation of *SOX18* observed upon their knockdown in HPAECs is particularly noteworthy. This consistency across multiple novel genes suggests that our PAH-former approach successfully identified candidates that influence a key PAH-associated cellular phenotype. In this way, fine-tuning Geneformer using the PAH dataset and subsequently conducting *in silico* perturbation analysis enabled us to identify a significant number of previously unknown disease-associated genes in PAH. Our approach provides a significant benefit over conventional DEG analysis by finding genes that causally affect cellular states, rather than simply indicating changes in expression levels. This ability to pinpoint functionally relevant genes, even from limited patient samples, demonstrates the unique utility of a transfer learning framework like PAH-former. While further validation is required, this approach offers a comprehensive means to explore therapeutic targets and has the potential to enhance the efficiency of fundamental experiments, reducing the effort and cost required for molecular function experiments. Moreover, the Geneformer and public database combination (fine-tuning) represents a promising new platform applicable to a wide range of diseases, particularly rare diseases where patient samples are scarce and the underlying pathological mechanisms are poorly understood. We believe it can greatly accelerate the advancement of our understanding of disease molecular mechanisms.

While our platform demonstrated promising capabilities, several limitations warrant consideration. First, the gene outputs are inherently dataset dependent. Different datasets, even those examining similar biological contexts, may exhibit variations in gene expression profiles, potentially leading to discrepancies in the identified key genes or pathways. Second, the fine-tuning process of our model is sensitive to the choice of fine-tuning datasets and hyperparameters, such as the learning rate. Variations in these parameters can lead to different model outputs and potentially affect the robustness of our findings. Third, our study’s focus on SOX18 mRNA expression as the primary validation metric is a limitation. While we have analyzed the expression levels of *SOX1*8, we have not yet validated whether these expression changes are directly linked to corresponding changes in cellular phenotypes relevant to PAH. Finally, we must acknowledge the multifactorial nature of PAH. Our gene-centric approach, while informative, may not fully capture the complexity of PAH, which likely involves intricate interactions of multiple genetic and environmental factors beyond single gene mutations, as well as cell-cell interaction change within the organs.

In conclusion, our novel Geneformer-based fine-tuning platform provides a powerful and broadly applicable strategy for disease-related gene discovery. This approach enables the identification and validation of new candidate genes, promising to advance cell-specific mechanistic insights and efficient therapeutic development for PAH.

## Methods

### Creating IPAH Dataset

Datasets utilized for fine-tuning and testing the Geneformer model were acquired from a publicly available database in the NCBI Gene Expression Omnibus (GEO). Specifically, we included datasets GSE169471 (Six control samples, three idiopathic pulmonary arterial hypertension [IPAH] samples), GSE210248 (three control samples, three IPAH samples), and GSE185479 (three control samples, three IPAH samples). Additionally, we incorporated a subset of the integrated Human Lung Cell Atlas (HLCA) v1.0 core dataset, specifically selecting samples annotated as lung parenchyma to augment the control dataset. Original datasets were obtained in ScanPy AnnData (h5ad) format. Quality control (QC) for GSE169471, GSE210248, and GSE185479 was conducted using the following criteria: total gene counts per cell ranging between 200 and 2500, and mitochondrial gene content below 5%. The HLCA dataset was used without further QC, as it was provided in a pre-processed format. For compatibility with Geneformer tokenization requirements, genes in the GSE210248 and GSE185479 datasets were annotated with their corresponding Ensembl IDs using the MyGene library. The GSE169471 and HLCA datasets already contained Ensembl IDs and were used directly without further modification. The prepared datasets were partitioned into three distinct groupings for downstream analysis:

A. GSE169471 only: four control samples and two IPAH samples for training, and two control samples and one IPAH sample for testing.
B. GSE169471 combined with HLCA: four control samples from GSE169471 plus 102 control samples from HLCA, and two IPAH samples for training; two control samples from GSE169471 plus five control samples from HLCA, and one IPAH sample for testing.
C. Combined datasets of GSE169471, GSE210248, GSE185479, and HLCA: For training, four control samples from GSE169471, two control samples each from GSE210248 and GSE185479, plus 102 control samples from HLCA; and two IPAH samples each from GSE169471, GSE210248, and GSE185479. For testing, two control samples from GSE169471, one control sample each from GSE210248 and GSE185479, plus five control samples from HLCA; and one IPAH sample each from GSE169471, GSE210248, and GSE185479.

### Fine-tuning of Geneformer

Fine-tuning was performed to classify PAH versus control cells by leveraging the Geneformer model pre-trained on extensive transcriptional data. Specifically, we obtained the gf-12L-95M-i4096 (12-layer Transformer block, 4,096-token maximum sequence length) model from the ctheodoris/Geneformer repository on the Hugging Face Hub. For implementation, we used PyTorch along with the Hugging Face Transformers library and executed training on an H100 GPU (NVIDIA). We adapted the pre-training setup, which initially employed a masked token prediction head, by replacing it with a sequence classification head suitable for the binary classification task (PAH vs. control). The fine-tuning hyperparameters were set as follows: a learning rate of 2×10^−5^, a batch size of 64, a cosine scheduler with 100 warmup steps, and a total of eight training epochs. To mitigate overfitting, the lower four Transformer layers remained frozen during training, thereby focusing updates on the upper layers while preserving the foundational representational capacity learned during pre-training. Three separate models were created, each corresponding to one of the dataset partitions (A, B, and C) described above. These models are referred to as model A, model B, and model C, respectively. Each model was fine-tuned independently using its respective training split and evaluated on the corresponding test set to assess its classification performance.

### *In silico* perturbation

*In silico* perturbation was conducted on models A, B, and C, following the approach described previously in the Geneformer study. Briefly, this method perturbed the gene expression ranking to simulate gene inhibition or activation within single-cell transcriptomes. Genes targeted for perturbation were comprehensively selected from those expressed in both control and PAH samples. *In silico* deletion was simulated by removing targeted genes from the rank encoding, measuring perturbation effects via cosine similarity changes in both cell-level and gene-level embeddings. Conversely, *in silico* overexpression was simulated by moving the targeted genes to the top of the rank encoding, modeling the activation of these genes. Perturbations were executed using the test splits of datasets corresponding to each model. Two scenarios were explored: perturbations transitioning from control to PAH states, and vice versa, each involving both *in silico* deletion and overexpression strategies. Genes exhibiting a false positive rate below 0.05 and demonstrating a decreased cosine similarity toward the target state upon perturbation were considered promising candidates.

### Enrichment analysis

To identify potential driver mechanisms underlying PAH pathogenesis, we applied Metascape ^36^ to genes extracted by our PAH-former. The results of each perturbation output by Geneformer were ranked by the cosine shift towards the goal state ("Shift_to_goal_end") (largest first) and the False Discovery Rate (FDR) (smallest first). We defined genes with a positive "Shift_to_goal_end" and an FDR < 0.05 as candidate disease-related genes.

Candidate genes whose *in silico* deletion shifted the cell state towards PAH were 134 genes in total and we used all of them for enrichment analysis. Candidate genes of other directions were more than 200 genes. For enrichment analysis, we used the top 200 genes for enrichment analysis.

### Target Gene Knockdown using RNA Interference

HPAECs were commercially obtained (PromoCell, C-12241) and handled according to the provider’s instructions. The cells were seeded in 96-well plates at a density of 2,400 cells/well in complete growth medium without antibiotics and incubated overnight. For *in vitro* knockdown experiment, the following siRNAs were purchased from Thermo Fisher Scientific: Silencer™ Select siRNAs for S100A6 (ID: s12418), TXNIP (ID: s12418), HSP90AA1 (ID: s6993), and MT2A (ID: s194629). siRNA was diluted in Opti-MEM I Reduced Serum Medium (Thermo Fisher Scientific, 31985070). For negative control, Silencer™ Select Negative Control No. 2 siRNA (Thermo Fisher Scientific, 4390846) stock solution was diluted with Opti-MEM I. Lipofectamine RNAiMAX reagent (Thermo Fisher Scientific, 13778150) was diluted with Opti-MEM. Equal volumes of diluted siRNA and diluted Lipofectamine RNAiMAX were combined and incubated at room temperature for 15 minutes to allow siRNA-Lipofectamine RNAiMAX complex formation. The volume of diluted Lipofectamine RNAiMAX solution was adjusted to use 0.2 µL of Lipofectamine RNAiMAX per well. The final concentration of each siRNA was 20 nM. Subsequently, siRNA-Lipofectamine RNAiMAX complexes were added to the cells. Cells were incubated with the complexes in a final volume of 120 µL per well at 37°C in a 5% COLJ incubator for 48 hours post-transfection.

Gene knockdown efficiency was evaluated at the indicated time points by quantitative PCR using QuantStudio 6 Flex Real-Time PCR System (Thermo Fisher Scientific). *SOX18* mRNA levels were evaluated for each condition at the same time. Data was normalized to the *GAPDH* expression level. The primer sequence used for the quantitative PCR analysis was as follows:

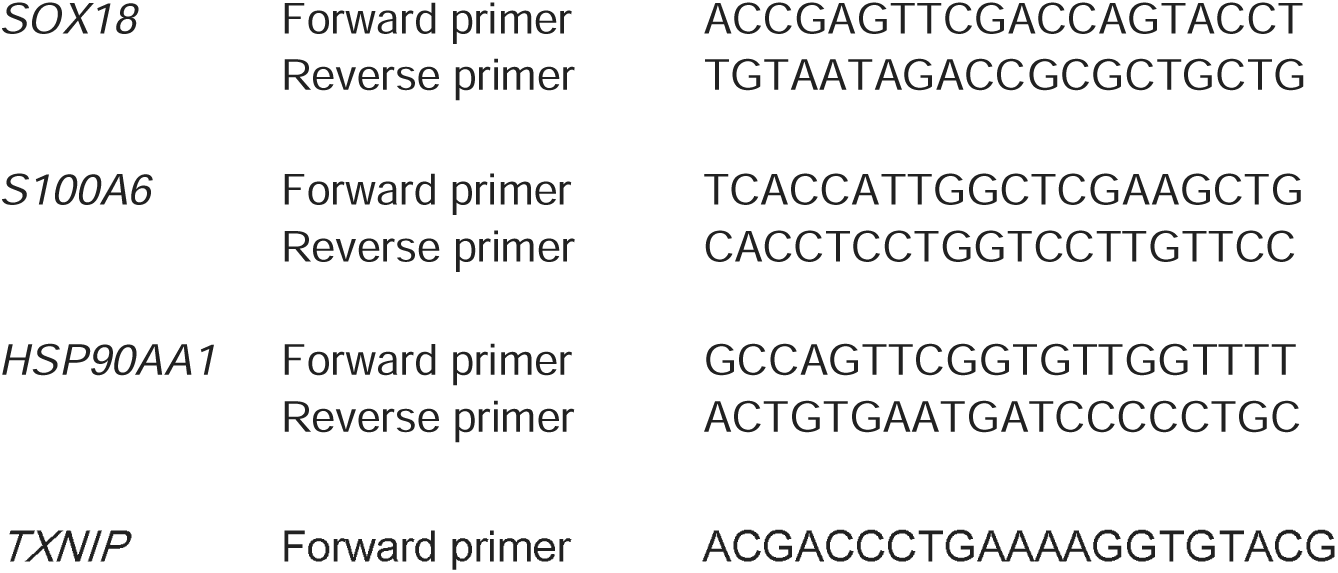
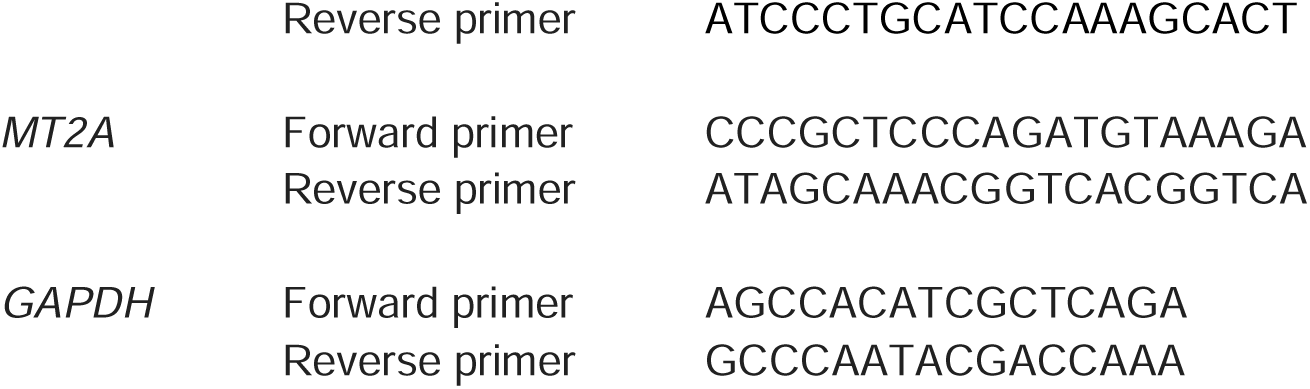

### Statistical information

All quantitative data are presented as mean ± standard deviation (SD). Statistical significance of differences between two groups was determined using a two-tailed Mann-Whitney *U* test. A *P* value of less than 0.05 was considered statistically significant.

Specifically, for the RNA interference experiments, the mRNA expression levels of target genes (*S100A6, HSP90AA1, TXNIP, MT2A*) and *SOX18* in HPAECs following siRNA-mediated knockdown were compared against control siRNA-treated cells.

For the knockdown efficiency assessment:

*S100A6* knockdown: Control (n = 9) vs. *S100A6* KD (n = 9), *P* < 0.0001. *HSP90AA1* knockdown: Control (n = 9) vs. *HSP90AA1* KD (n = 9), *P* < 0.0001. *TXNIP* knockdown: Control (n = 9) vs. *TXNIP* KD (n = 9), *P* < 0.0001.

*MT2A* knockdown: Control (n = 9) vs. *MT2A* KD (n = 9), *P* < 0.0001.

For the assessment of *SOX18* mRNA expression changes following knockdown of candidate genes:

*S100A6* knockdown: Control (n = 3) vs. *S100A6* KD (n = 8), P = 0.0485. *HSP90AA1* knockdown: Control (n = 3) vs. *HSP90AA1* KD (n = 9), *P* = 0.0091. *TXNIP* knockdown: Control (n = 3) vs. *TXNIP* KD (n = 9), *P* = 0.0091.

*MT2A* knockdown: Control (n = 3) vs. *MT2A* KD (n = 8), *P* = 0.1939.

All statistical analyses were performed using GraphPad Prism version 10.4.2 (Dotmatics).

### Data availability

The datasets generated and analyzed during the current study are available in the Figshare repository.

### Code availability

The pretrained PAH-former models, including those trained on three distinct datasets, are publicly available via the Hugging Face Hub repository (https://huggingface.co/so298/PAH-former). Each model is published on a separate branch: "base_dataset," "add_hlca," and "add_other_data." Additionally, the source code for the PAH-former analyses is openly accessible through our GitHub repository (https://github.com/UTcardiology/PAH-former-analysis).

## Acknowledgements

This work was supported by Cross-ministerial Strategic Innovation Promotion Program (SIP) on “Integrated Health Care System” Grant Number JPJ012425. We would like to thank Yukiko Kaneko for her technical assistance.

## Author contributions

T.K. and S.H. designed the study, interpreted the results, and wrote the manuscript.

M.I. conceived the study and supervised the project.

S.H. developed the computational model, performed in silico perturbation analyses.

T.K. designed and performed the RNA interference experiments, analyzed the experimental data.

S.K. provided the computational resources necessary for the AI model and supervised the project.

A.K., R.T., S. M., J.I., T.I. and N.T. supervised the project. All authors reviewed and approved the final manuscript.

